# Global worming: massive invasion of North America by earthworms revealed

**DOI:** 10.1101/2022.06.27.497722

**Authors:** Jérôme Mathieu, John Warren Reynolds, Carlos Fragoso, Elizabeth Hadly

## Abstract

Human activities cause major ecological changes by reshuffling the spatial distribution of species. The extent to which this process affects belowground biota is a critical issue because soil organisms play a key role in ecosystem functioning and maintenance. However, the magnitude of the reshuffling of soil species remains unknown so far because of the lack of a historic baseline. Here, we begin to fill this gap with the largest spatiotemporal database of native and alien earthworms in North America. Our results reveal that the entire continent is being invaded by non-native earthworms through a variety of pathways. We show that these aliens bring novel ecological functions in most regions and thus represent a major threat to native ecosystems. Our findings demonstrate that earthworms, and most likely other soil organisms, represent a major but overlooked pool of invasive species with strong ecological impact. They need to be better integrated in control and mitigation strategies.

## Main Text

The transport of species beyond their biogeographic barriers plays a major role in global homogenization of biodiversity^1^, biodiversity loss, and has countless consequences on ecosystems (*1*). Invasive insects alone cost a minimum of US$70.0 billion per year globally^2^. Biological invasions are a major driver of global change, synergized by and influencing biodiversity loss, climate change and land use change^3^. Identifying aliens that can turn invasive and anticipating their spread is thus critical to build environmental management strategies and policies.

The spatial distribution and dynamics of alien species is now well documented at the global scale in a large number above-ground taxa^4^. This knowledge provides a background to develop management strategies of aliens at national and international scales. In contrast, the spatial distribution of alien species belowground is still largely unknown^5^. A limited number of studies documented the expansion of a few specific soil organisms, and to date, none have quantified the colonization dynamics of multiple soil species on a continental scale. Ignoring these organisms will result in missing critical mechanisms of ecological changes for soil organisms have a strong impact on overall ecosystem function. Many soil organisms are qualified as ecosystem engineers^6^ precisely because they play a central role in many processes that cascade to aboveground communities and to the atmosphere^7^. Consequently, any differences in life histories, network linkages (i.e., trophic or symbiotic), and metabolisms of alien soil species will have a strong impact on native ecosystems.

Among alien soil taxa, earthworms are of particular concern because of their dramatic impact on native ecosystems. Earthworms, in general, are viewed favourably because they are perceived to increase crop productivity and maintain soil fertility^8, 9^. They are iconic of good soil ‘management’. As a consequence, colonization by alien earthworms is often viewed positively, and sometimes even encouraged^10–13^. However, not all earthworms are the same. In tropical areas, alien earthworm species such as *Pontoscolex corethrurus* can result in compacted soil and make it unsuitable for plant growth in less than a year^14^. In the northern broadleaf forests of USA and Canada, particularly where few earthworms survived Pleistocene glaciation, colonization by alien earthworms affects all components of the native ecosystems^15^. Native soils are transformed by the mixing of the horizons and the rapid disappearance of the litter layer^16^. Alien earthworm bioturbation results in a simplification of the understory vegetation and increases mortality of important economic species such as sugar maple (*Acer sacharrum*)^17–19^. Alien earthworms also impact native soil food webs and the organisms that depend on them, such as springtails, ground beetles and the red-backed salamander^20, 21^. Ultimately, the colonization by alien earthworms can facilitate the spread of invasive plants^22^.

Even though the strong effects of alien earthworms are well known, their spatial distribution and their colonization dynamics over large scales has not been well characterized. Therefore, we have limited baselines to assess where alien earthworms may represent a threat, or where they have already impacted ecosystems. This is surprising because it is known since the beginning of the 20^th^ century that several species of earthworm have colonized different parts of the world by inadvertent human transport^23^. Indeed, it was predicted in 1900 that alien earthworms should have conquered the whole of North America by the end of the XX^th^ century^24^. But, to date, no data were available to test this prediction. North America is particularly sensitive to biological invasions^25, 26^. Many North American ecosystems evolved in the absence of earthworms at least since the Last Glacial Maximum, until their recent introduction by humans ^27^. In these ecosystems, earthworms arrived into an empty belowground niche with limited competitors. In addition, climate change is facilitating their spread from South America to Mexico and into the northern parts of the continent, where the permafrost is melting, potentially introducing a positive feedback loop with climate change^28–30^.

Here, we reconstruct the spatiotemporal spread and putative introduction pathways of exotic earthworm species at the scale of a continent over a century. To achieve this, we built the database EWINA, which is the most comprehensive database of native and exotic earthworm species occurrence ever built. This database collates more than 68k records dating from 1850 to 2021, across 2510 geographical units in North America (Extended Data Fig. 1). We completed this database using EWINA_IPATHS, a second database which focuses on the introduction pathways of alien earthworm species into the USA between 1945 and 1975. Combined, these two databases provide a baseline with which to ascertain earthworm invasion dynamics in North America, which can be used to investigate alternative scenarios and policies for management of alien earthworms and other soil taxa here and globally.

## Results

### A massive invasion of North American soils by earthworms

To examine the extent of the geographic distribution of alien earthworm species, we mapped the predicted Relative Alien Species Richness (RASR) of earthworm across North America (Fig. 1A), based on present environmental condidtions. RASR is the ratio of the number of alien species over the total number of species in a given geographical unit (typically a county herein, see Extended Data Fig. 1). We modeled RASR as a function of 12 environmental drivers (Extended Data Table 1) using data between 2000 and 2021, with a machine learning approach. Predicted RSAR of earthworm in actual conditions is overall extremely high – with a median of 73% across North America (Fig. 1B), meaning that alien species dominate in a major proportion of the continent. Only 3% of studied geographical units are devoid of alien earthworms, while 28% are devoid of any native ones. The map reveals an opposition between the northern part of the continent, which has a RASR typically higher than 50%, and the south and west of the continent, where RASR is typically below 50%.

**Figure 1.**
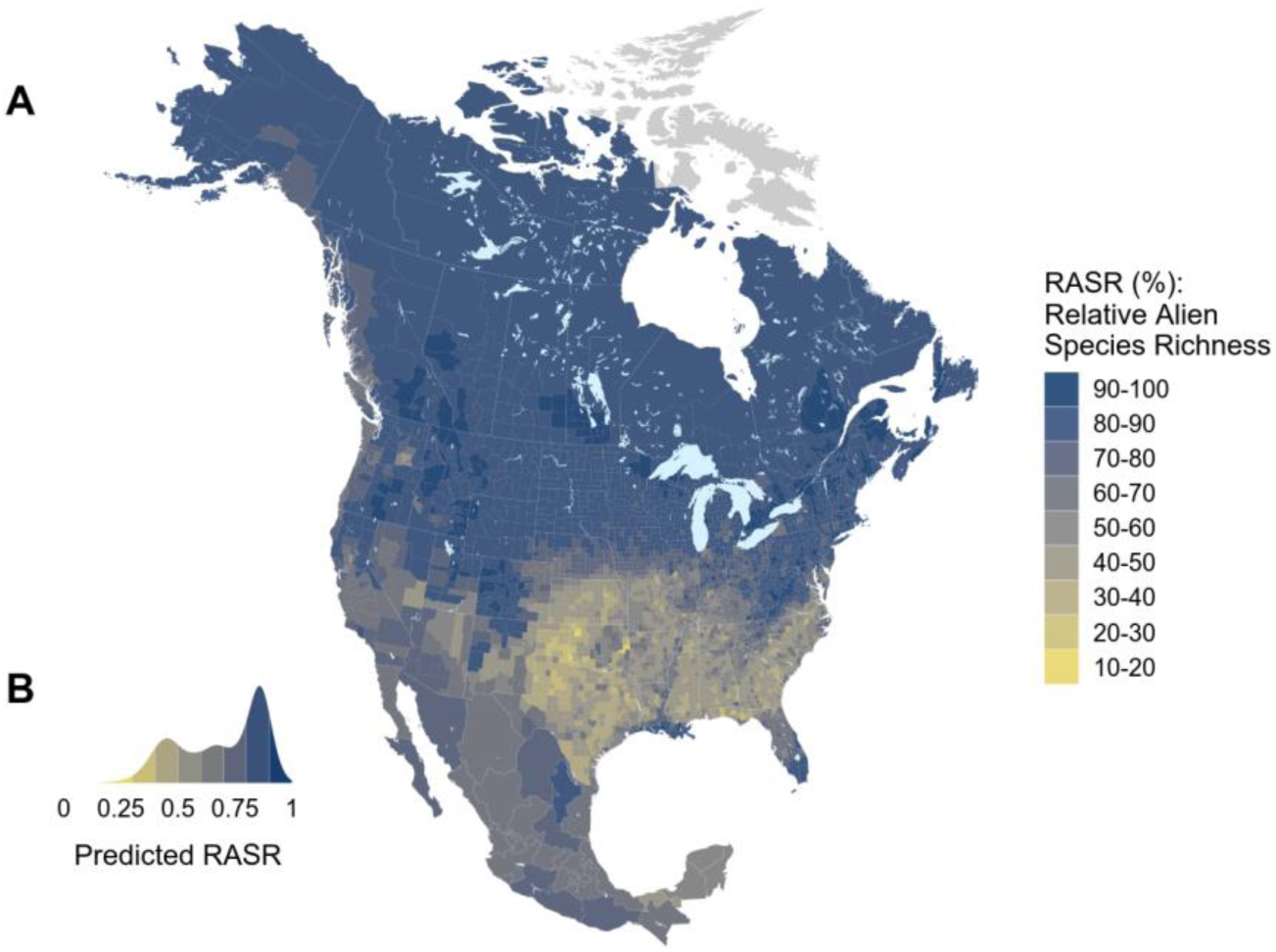
Relative alien earthworm species richness in North America. (A) Map of predicted Relative Alien Species Richness of Earthworm in North America (RASR). The colors indicate the value of predicted RASR in each geographical units. RASR is calculated as the proportion of species within a geographical unit that are alien. (B) Statistical distribution of the predicted values of RASR across the geographical units.

**Table 1.**
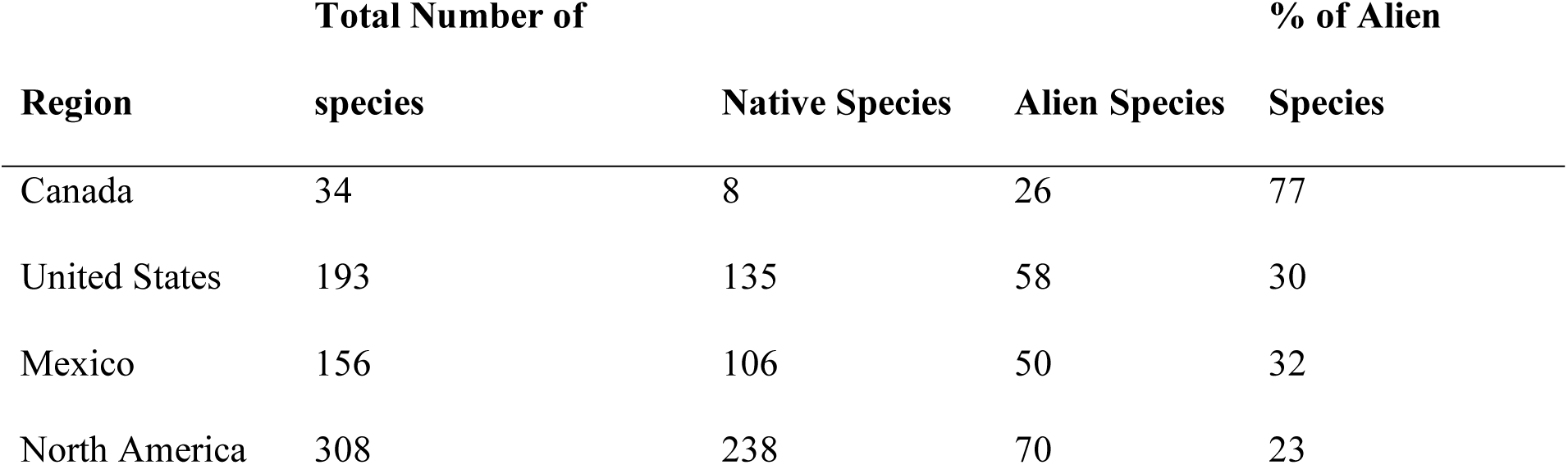
Number of native and alien earthworm species in North America.

Overall, exotics represent 23% of the 308 earthworm species recorded in North America (Table 1). RASR at the country level is twice as high in Canada than in the US and in Mexico. There are many more alien species than native ones in Canada. Over North America, exotic earthworms have on average a larger geographical range than the natives (Extended Data Fig. 2), typical of other invasive species. A startling twelve of the thirteen topmost widespread earthworm species are alien. *Aporrectodea trapezoides* and *Lumbricus rubellus* are the most ubiquitous species (60 and 43% of geographical units respectively). Alien species are also proportionally more parthenogenetic than native ones (Extended Data Fig. 3).

### Climate is the best predictor of aliens’ distribution

Surprisingly, climate, but not human activity, was the strongest predictor of RASR (Extended Data Fig. 4). Mean Annual Temperature (bio1) clearly had the highest effect on RASR, followed by carbon at soil surface and annual precipitation (bio12). The other environmental features, including human activity, had a very limited impact on RASR. Mean annual temperature and annual precipitation were negatively related to RASR (Extended Data Fig. 5). Soil carbon at soil surface was positively related to RASR. The effect of the other covariates was very limited, except the human impact index, which was positively related to RASR in a few environmental conditions (Extended Data Fig. 5).

### Earthworm alien species richness is correlated to above ground taxa

To test the specificity of the distribution of alien earthworms versus aboveground exotics, we correlated the alien earthworm species richness to the one of non-native Plants, Spiders, Mammals, and Birds across the geographical units defined by the Biodiversity Information Standards^31^ (TDWG). We used the data from Dawson et al.^4^ to compute aboveground alien species richness (Extended Data Fig. 6). Earthworm alien species richness was positively correlated to all taxa, particularly with plants and spiders (Fig. 2). This suggests that alien earthworm species richness in North America is not independent to that of aboveground taxa and indeed, they may be linked, at least at the resolution of the TDWG units.

**Figure 2.**
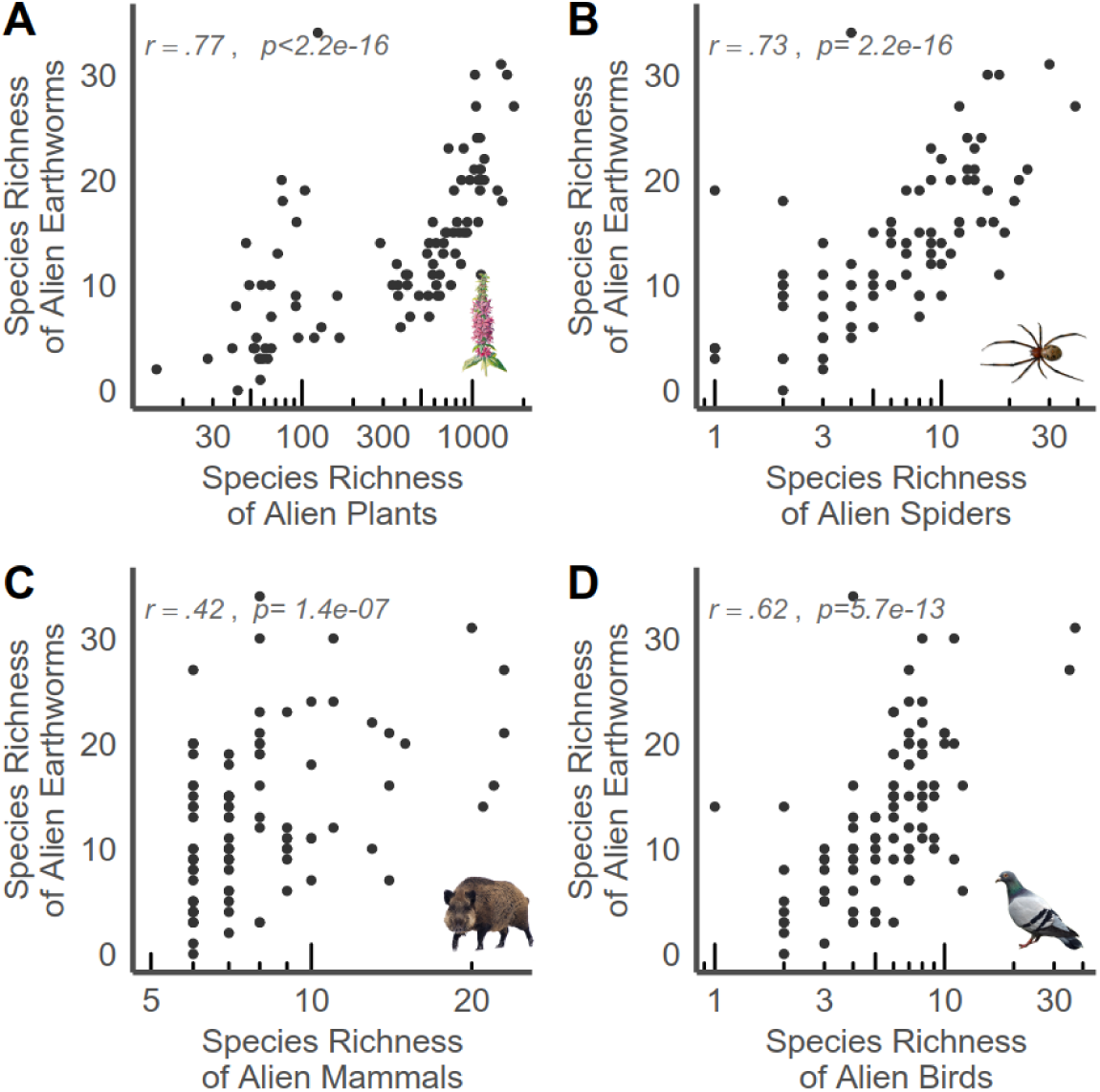
**Correlation between earthworms and aboveground groups alien species richness**. Each dot represent the number of Alien Earthworm Species Richness and (A) Plants, (B) Spiders, (C) Mammals, (D) Birds across TDWG level-4 regions in North America. Data of these groups come from Dawson et al.^4^. The correlations were tested with the Spearman correlation test.

### A slow but certain colonization of North America from Europe and Asia

Anticipating the spread of alien species requires a general understanding of the temporal dynamics of the colonization process. To tackle this issue, we evaluated three critical aspects of the dynamics.

First, we looked at how alien species number have developed in time in North America since 1860 (Fig. 3A). The temporal accumulation of alien species is constant and steep after 1900, with a peak of first species records in 1950. The average rate of colonization since 1950 is one new alien species every three years.

**Figure 3.**
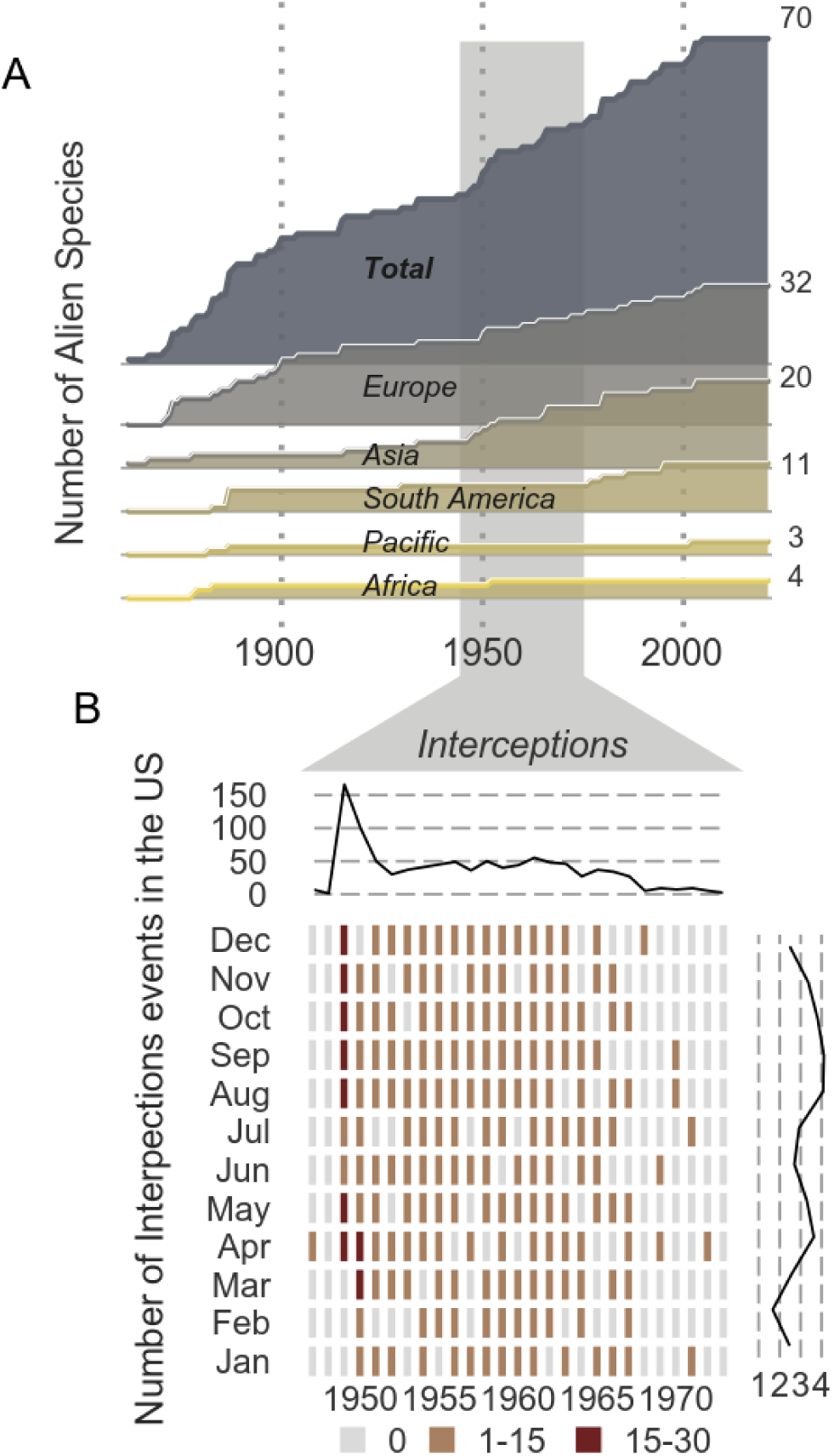
Historical dynamics of North America’s colonization by alien earthworm species. (A) Temporal Evolution of the number of alien earthworm Species recorded in North America (Mexico, United States, and Canada). The number of established species is detailed by the continent of origin of the species. The respective total number of alien species are indicated separately on the right. (B) Temporal trend of interception rate of earthworm specimens at the US borders between 1945 and 1975. The continuous line on top indicates the number of interceptions per year. The continuous line on the right indicates the average monthly number of interceptions. The color of the tiles indicates the monthly rate of earthworms intercepted at the US Borders.

Second, we assessed the strength of the propagule pressure - the flux of individuals and earthworm species - arriving from other places of the world. To do so, we compiled reports of events of earthworm interceptions at the US borders over 30 years. Although these data are not standardized and only represent a fraction of the total introductions, they show that introductions were happening consistently between 1945 and 1970, with a peak in 1949 (Fig. 3B). This means that apart from 1950, the propagule pressure was fairly constant over this period of time.

Third, we analyzed the nature of the introduction pathways and their geographical features. Alien earthworm species principally originate from Europe and Asia (Fig. 4A). Most species were introduced by airplane (Fig. 4B). Islands such as Hawaii and Caribbean islands seem to have played an important role as both sinks and sources of alien species. Alien earthworms were mostly intercepted in coastal regions of the US, in airports and harbors, which is consistent with the observed spatiotemporal colonization pattern (Fig. 5).

**Figure 4.**
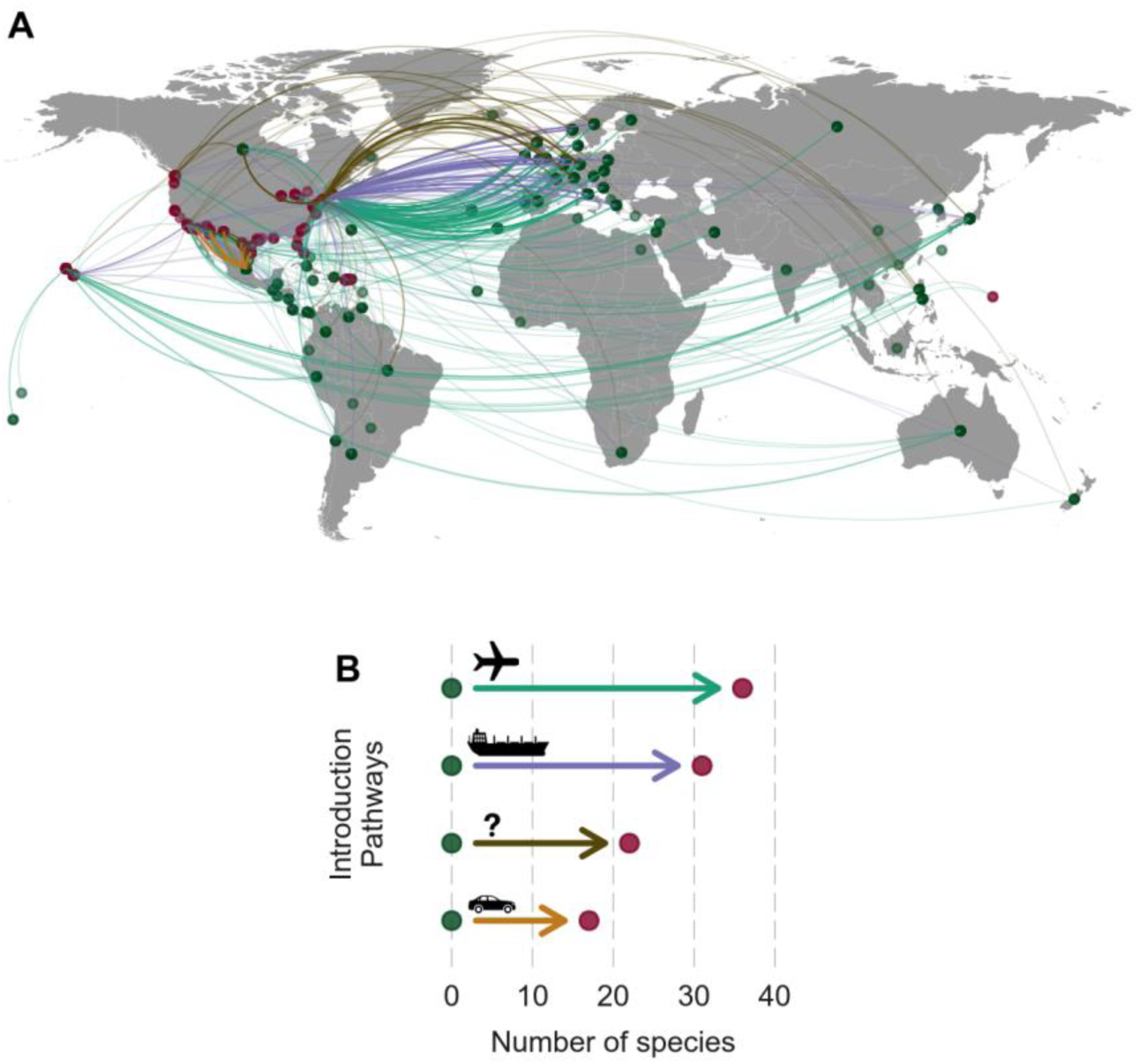
Introduction pathways of alien earthworm species in the United States between 1945 and 1975. (A) Each line indicates an intercepted introduction pathway of earthworms at the US border. The green dots indicate the origin of departure, while the red ones indicate the location of interception in the United States. When the exact location of departure was not known, we located it in the center of the country of departure. The color of the lines indicates the transportation means, following (B): The number of species intercepted through the different introduction pathways. Please note that Puerto Rico and Guam islands were part of the United States over this period.

**Figure 5.**
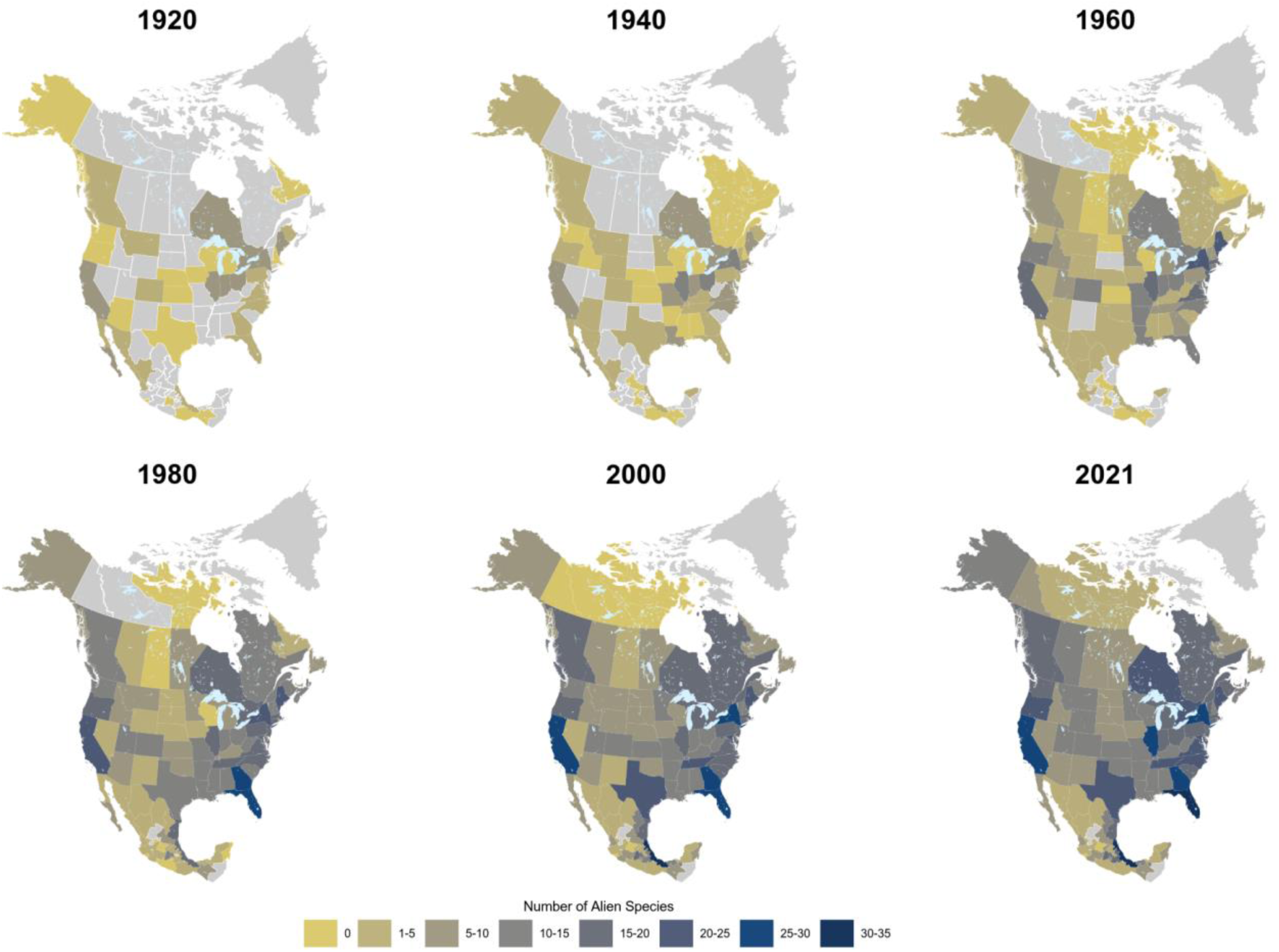
Spatio-temporal dynamics of earthworm alien species richness across North America. These maps relates the accumulated number of known alien earthworm species in TDWG4 units across North America at different dates. The colors are proportional to the number of exotic species. Zero means that only native species have been reported at the considered date. Gray colors represents areas with no data.

### A colonization from the coast to the interior

To determine the colonization rate of aliens within the continent, we reconstructed the spatial temporal colonization of alien earthworm species across North America since the beginning of the 20^th^ century (Fig. 5). Even though data are scare before 1960, it appears that colonization started from the both Atlantic and Pacific coasts - where major ports were located - and gradually reached the interior of the continent. The temporal dynamics of the individual alien species spread reveals a continuous and pulsed increase in the geographical range of species (Extended Data Fig. 7).

### Alien earthworm species bring ecological novelty

To assess the potential impact of alien earthworm species on native ecosystems, we evaluated their ecological redundancy with native earthworm species. High redundancy would suggest little potential impact, while low redundancy would suggest ecological novelty and high potential impact. We mapped the functional enrichment brought by the arrival of alien earthworm species across the TDWG units to identify the zones under highest threat. Native and alien earthworm species share little functional redundancy (Fig. 6A). Native species mostly feed in/on soil, while aliens primarily feed in/on litter. This suggests that in the majority of locations, the arrival of alien earthworm species at least increases litter decomposition. In locations where no native earthworms were present, such as the northern Great Plains, aliens should also increase soil bioturbation. Overall, alien earthworms bring higher rates of biomass turnover except in the parts of the eastern USA and a few locations in Mexico and the west coast.

**Figure 6.**
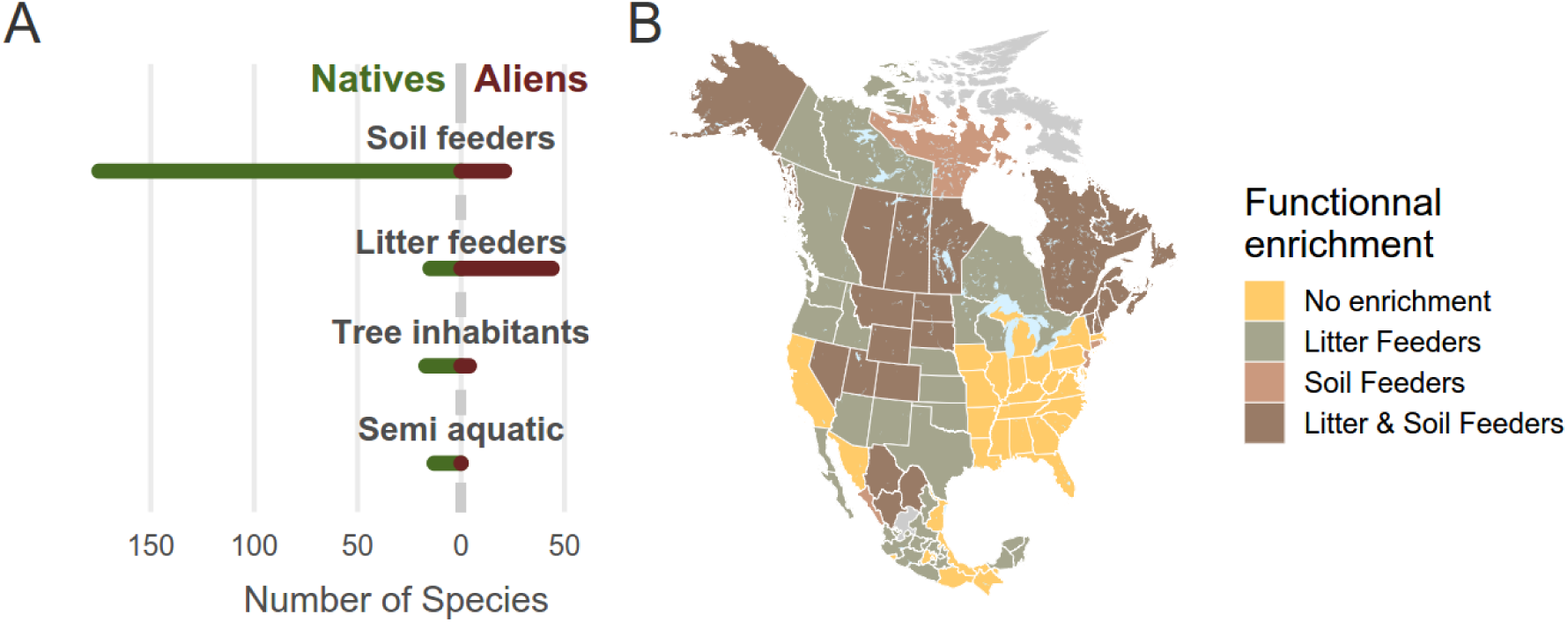
Functional niche of earthworm species in North America and functional enrichment brought by aliens. (A) Functional niche of Native and Exotic species. The bars represent the number of species that feed or inhabit a specific food or habitat. A species can occur in several categories. (B) Ecological function brought by alien earthworm species.

## Discussion

### A massive colonization of North America by alien earthworm species

The majority of the North American soils contain alien earthworms. Only 3% of the geographical units that we investigated were free of exotic earthworm species. At the county level, RASR was dramatically high, with a median above 70%, showing that in most of the continent, alien earthworm species are in fact more diverse than the native species. Temporal dynamics suggest that there may be a limit to the introduction of new alien earthworm species, at least in the present environmental conditions. However, the spatiotemporal maps suggest that alien species are still becoming invasive within the continent. Overall, our study demonstrates that multiple species of earthworms, and by inference, other soil macro and microinvertebrates, have been massively translocated between and within continents, and should be considered in biological invasion policies where they are not already. For example the trade of alien earthworm species is allowed without restrictions between most states of the US.

It is difficult to assess whether the trend we found is specific to North America because of the lack of other similar databases on alien earthworms in other parts of the world. The few other available data on earthworm alien richness suggest that they may also be substantial in other regions. For example, the Northern Russian Plain is believed to be inhabited mainly by European alien species^32^. A few countries such as China, Korea, and Myanmar also seem to host a high level of alien species, between 20 and 30%^33^. At the other end of the gradient, Africa is believed to have very few alien earthworm species^34^, though no quantitative data are available to confirm these statements.

Surprisingly, earthworms’ RASR increases from the tropics to the North Pole, whereas it decreases towards the pole in other aboveground terrestrial invertebrates^35^. In contrast, alien species richness of earthworms shows a parallel gradient with some aboveground invasives, especially plants and spiders. Correlation with plants and spiders is likely due to a common history of introduction, as earthworms and spiders are often imported accidentally as contaminants of plants, either in the soil around the roots, in the leaves, or among fruits^36^.

Earthworms are easily at the top of all invasive animal taxa in North American in terms of alien versus native species richness ratio, with 25% of North American earthworm species being exotic. In the United States alone, only 6% of mammal species are exotic, 2% of insects and arachnids, and 8% of fish species^37^. The lack of awareness of how much the soil biota has been transformed is alarming. The high proportion of alien species in the earthworm group may be explained by the limited pool of North American native species: 240 species, likely due to the absence of native earthworms in much of the continent for reasons that are still debated^27, 38^. The presence of the Ice Sheet during Last Glacial Maximum (LGM) in North America has often been put forward as an explanation of this peculiar pattern. Another – nonexclusive - explanation is that the diversity of native earthworm species is underestimated because not all native species have been described, particularly in southern and central parts of the continent.

Accidental introductions are typical of invertebrates in general^39^. We show that this dynamic in earthworms was sustained over 30 years and that the species originated mostly from Europe and Asia. Interestingly, the peak of new aliens in 1950 was also reported in insects and mollusks ^40^. It can be linked to the increased trade that followed the General Agreement on Tariffs and Trade (TAGG) of 1947, which reduced trade barriers such as tariffs and quotas. Overall, our results suggest a strong link with the trading routes and policies. It must be noticed that our data do not cover legal importations, in particular between Canada and the US. Indeed, over 500 million alien earthworm now are exported yearly from Canada to other countries, especially to the USA, to be used as fishing or composting commodity^41^. This massive flux constitutes a major and legal pathway of alien earthworm species transportation. It appears in contradiction with the efforts developed by the U.S. Department of Agriculture to prevent introductions of alien earthworms in the US.

### Invasive or alien earthworms?

Due to their large distribution and their specific ecological role, alien earthworms represent a serious threat to native ecosystems in North America. Whether these organisms will turn into invasive depends on their impact on native ecosystems. The impact of alien earthworms in broadleaf forest has been well documented, showing that these ecosystems are under particular threat. However the impact of alien earthworms on other native ecosystems such as natural grasslands, conifer forests, or chaparral, characteristic of unique species assemblages in North America, or in our croplands, is poorly known (but see^42–45)^. It may be more critical to deliberate on the status of alien earthworm in these systems. We show that when alien earthworms colonize a new area, most of the time they fill an empty functional niche (Fig. 6), because natives are absent or tend to be soil feeders, while aliens tend to feed on litter. Aliens that establish thus have a strong potential impact on native ecosystems.

It is thus tempting to classify non-native earthworm species as invaders. However, many alien earthworm species in North America provide important ecosystem services, particularly to agriculture^7^. For instance, their presence dramatically increases crop productivity^7^, which motived their deliberated introductions in many places of the world^10, 11^. In some parts of Canada, alien earthworm picking and trading represent an important economy^41^, demonstrating that although alien earthworm species may be considered as a pest by one sector, they may be a valuable ecosystem provider to another^46, 47^. Earthworms highlight that the one major group of organisms may be perceived as beneficial in some regions, typically agricultural ones, but negative in others, such as more natural ones, a case similarly derived for the European honeybee^48^. We know little about how either native or non-native earthworms have impacted agricultural lands or less-impacted ecosystems.

### Earthworm’s spread in the future

Expansion of earthworms in North America will depend both on the flux of introductions into the continent and on the capacity of aliens to spread within the continent upon arrival. The temporal trend in the accumulated number of alien earthworm species shows a plateau since 2001. This plateau suggests that the number of earthworm species in North American soils may have stabilized. But this doesn’t mean that the corridors for new species or for in situ increase in abundances have vanished. For example the flux of aliens between Canada and the US is still massive due to the fishing bait market^41^, and the further demise of native earthworms may induce new limits to introductions and invasions.

The spatial spread of alien species within the continent directly depends on the dispersal capacity of earthworms. Dispersal of earthworms is mostly passive and achieved by the transport of individuals and human activities^49^. In North America, known transports include uses for fishing, vermicomposting, and agriculture. It has been predicted that in a business as usual scenario, 49% of suitable habitats in northern Alberta will host alien earthworm species within 50 years, and more than 92% in Ottawa within a century^50, 51^. However, there are large uncertainties around these estimations because the population dynamics of these species, and their competitive interactions with natives, are poorly known. Our results suggest that aliens will increase in numbers and distribution rapidly because many are parthenogenetic (Extended Data Fig. 3). However, we need a better understanding of life histories of both native and alien species to be able to anticipate their potential development at local-to-continental scales.

Our results also suggest that climate change will play an important role in the future spread of alien earthworms, because temperature and precipitation are among the most important predictors of earthworms’ RASR. Climate has also been identified as the main driver of future biological invasions^52^ in other invertebrate taxa such as ants^53^ and gastropods^54^. Though, whereas invasion hotspots of ants and several other taxa are predicted to occur in warm areas^53, 55^, alien earthworms are more prevalent in colder, wetter areas. Hence, alien earthworm species may expand northwards more rapidly to track cool, humid soils, which are more favorable to the exotic species. Canada and northern parts of the US thus appear as potential future hotspots for alien earthworm species in changing climates^56^. In southern regions, climate change will less likely facilitate the spread of alien earthworms from Asia and Europe. However, warming of temperate areas may open colonization opportunities for tropical alien earthworm species, as already observed in a few temperate grasslands^57^.

Because it is virtually impossible to remove established populations of alien earthworms^58–62^, the best management options we have is to focus on prevention and early detection^63^. Prevention can take several forms such as education of gardeners, fishermen and farmers. Such effort has already begun in the regions of Great Lakes ("Great Lakes Worm Watch" initiative) and could be extended via citizen science, through for example the #WorldWormWeek. Another approach is to encourage scientific research and use of native species in fishing and vermicomposting of agriculture. For instance, alien composting species such as *Eisenia foetida* can be replaced by native ones such as *Bimastos tumidus* ^64^. Several species of the native genus *Diplocardia* are currently collected commercially for bait from natural populations in Kansas, Missouri, and Florida^64^. Overall, because earthworms are dispersal limited but easily transported by human activities, the future spread of alien earthworm, and probably other soil organisms, strongly depend on our capacity to develop policies to manage fluxes of alien soil organisms at national and international levels^62^.

## Supporting information

Supplementary file 1 - list of references

## Methods

We built two separate databases: EWINA_RICH, which gathers native and alien earthworm species richness across North America, and EWINA_IPATHS, which collates data of earthworm interceptions at the US Borders.

### EWINA_RICH: Native and alien Species richness

To estimate the magnitude of introductions of Alien earthworm across North America, we calculated the Relative Alien Species Richness (RASR) of each of the North American geographical units, as defined below. RASR was computed as the ratio of the number of alien species versus the total number of earthworm species (natives and aliens). This metric has numerous advantages: it is independent of region size and scale of analysis, it is comparable across regions and ecosystems and between groups of taxa^65^. We used only data from 2000 to 2021 for the modeling and mapping of RASR.

To build the database, we compiled data of both native and alien earthworm species occurrence across 2510 geographic units in North America (Mexico, United States, and Canada, Greenland), covering 73% of the land surface (Extended Data Fig. 1). The data gathers the species name, the locality and date of observation, and when available, the abundance, the geographical coordinates and habitat features. The database contains individual records of species, but also community data, where all species were sampled together at the same place. We used the finest possible geographical resolution to define the geographic units, which in the US and Canada is the level 3 of the GADM (https://www.gadm.org). It corresponds to counties in the United States and to Parishes in Canada. The earthworm data were collated from 459 sources dating from 1891 to 2021. Data were searched in several manners. First, we compiled all data published or edited by the authors. J.W Reynolds (JWR) published over 500 articles and 25 books about earthworm’s spatial distribution in North America. He is the chief editor of the Journal Megadrilogica since 1972. This journal specializes on the ecology of earthworm in North America. JWR regularly published accounts of earthworm distribution in North America. As a senior author and editor, he has a deep knowledge of the literature published regarding earthworms in North American.

Carlos Fragoso worked over 35 years on the distribution of earthworms of Mexico and published over 75 articles and 46 book chapters, with a specific emphasis on native and exotic species. He published the great majority of available data regarding Mexico. In a second step, we reviewed systematically all the 391 articles published in the Journal Megadrilogica. We identified 230 papers from this journal with earthworm data and extracted the data from the text and from the maps. After this step, we compiled all references cited in these papers, get all possible original sources, and entered their data. To complete this process, we compiled the list of species present in North America and did a specific search for each native species. Once this step was achieved, we established the list of authors who published data on native earthworm in North America and did a systematic search of all the papers published by these authors. Finally, we did classical research on web of science and Google Scholar, with two approaches. We combined the key word "earthworm" or "oligochaete" and a key word for location. We used several location key words such as country names and each state or province names of North America. After this step, we used the list of all native species to do a search on web of science and in GBIF. In order to minimize identification issues in GBIF, we removed all GBIF data originating from human observations and kept only the trusted ones from museums and molecular biology portals. Once all data were compiled, we homogenized the taxonomy with the Csuzdi’s database^66^ and the GBIF Backbone Taxonomy. We aggregated subspecies at the species level. We obtained 68 938 records, from which we eliminated the duplicates based on their locality and date of observation (month and year). At the end, the database holds 66 512 unique species occurrence. With this list of species occurrences, we computed the list of species by geographical units at two resolutions: at county level and at TDWG-level 4 units. TDWG4 units are standardized geographical entities used to compare biodiversity data across taxa^31^. Counties boundaries were obtained from GADM level 2 (parishes) in Canada, from the tiger 2014 database of the US Bureau of Census for the US, and from GDAM level 1 for Mexico. The TDWG-4 boundaries were obtained from a paper^4^ on invasive species from which we extracted data (see below). The different spatial layers were then merged with snapping activated and cleaned by geo-scripting to fix geometry issues such as duplicated points, auto intersections or sliver polygons. Last, we checked and fixed topology by neighborhood analysis and simplified the boundaries of the polygons, while preserving topology, with mapshaper.

### EWINA_IPATHS: Introductions pathways’ database

The second database, "EWINA_IPATHS" (Earthworm Introduction Pathways) documents the introduction pathways of earthworm in the United States. This database centralizes data of earthworm interception events at the US borders between 1945 and 1975. These data came from the U.S. Bureau of Plant Quarantine, U.S. Department of Agriculture. They were principally reported by G.E. Gates in a list of articles published between 1956 and 1982. Each record in the EWINA_IPATHS database relates an interception event of introduced earthworms. Interception events are described by the name of the intercepted species, its abundance, the date and locality of interception, the geographical point of origin, the transportation mode (boat, plane, car), and the substrate in which the earthworms were found (e.g. soil, leaves, fish bait). EWINA_IPATHS contains 1016 events of earthworm interceptions.

### Definition of alien and native species

The geographical origin of earthworm species is relatively well established at the country level in North America. Historically, due to the absence of earthworm fossils, the origin of species was estimated from the distribution of group of species and genus. Indeed, it is well known that certain genus such as *Lavellodrilus* are restricted to certain regions of North America and are considered native for this reason. More recent work based on molecular biology revisited these first imputations. They confirmed that in general alien species belong to distinct taxonomic and phylogenetic groups^67^. For our analyses, we considered the native/alien status of each species both at the continent and at the country scales. At the continent scale, we considered species as native when they were native in any of the three countries of North America. This classification was used for the global analyses of the North American continent, such as the cumulative number of alien species (Fig. 3) or the analysis of species attributes such as functional type (Fig. 6A) , reproduction (Fig.S3) and spread dynamics (Extended Data Fig. 7). A few north American species, for example from the genus *Ramiellona* and *Arctiostrotus,* were native from only one or two North American countries and could be considered as alien in other parts of North America. This was the case for species that were rare outside a clearly delimited area and were only present in disturbed areas outside of these regions. To take into account this fact, we also defined the native/alien status of each species at the country level. This system was used for the analyses at the TDWG4 levels or county levels (Fig 1, 2, 5, 6B, Extended Data Fig. 6).

### Environmental drivers of RASR

We tested 12 environmental features as potential determinants of RASR level (see Extended Data Table 1 for a full description of all examined variables).

First, we selected variables that were identified as global drivers of invasions in several other aboveground groups. These general macroecological variables included climate^68^ : Mean Annual Temperature (bio1) and Annual Precipitation (bio12), human activity (Global Human Influence Index^69^), the proportion of grassland and cropland in landscape^70^, and two habitat features: elevation and elevation heterogeneity (GTOPO30^71^).

Second, we added environmental features that are - according to previous studies - specific to earthworm invasions ^50, 72^. These variables were related to soil properties: the soil carbon content on surface and at 30 cm depth, the soil pH in the ten first centimeters^73^, as well as the (log) density of roads^50^ (length of roads by units of surface). We included the effect of the area of geographical unit in order to estimate any bias associated to the size of the geographical units, but this variable had no effect (Extended Data Fig. 4 and 5). Finally, we checked multi- collinearity among predictors using qr-matrix decomposition and pairwise plots. No covariates were correlated.

### Model of RASR

We modeled and predicted RASR as a function of the 12 environmental covariates with a random forest model. We used only observations posterior to 2000 and removed geographical units with less than five species, to remove under sampled units. The model was bagged in order to assess the test error without the need to perform additional cross validation to estimate the predictive accuracy^74^. Bagging (Bootstrap aggregation) consists in drawing many random samples from the training dataset (bootstraps), fit a separate model on each bootstrapped dataset, and average the predictions. On average each model use around two-third of the observations.

The remaining one third observations, not used to fit the model, are referred to as the out of bag (OOB) observations. We can predict RASR and estimate the prediction accuracy of each observation using all the models in which the observation was OOB. The prediction is obtained by averaging these predicted responses. With this approach, the out of the bag error is a valid estimate of the test error for the bagged model^74^.

The performance of the model was checked by plotting the predicted RASR versus the observed RASR (Supplementary Extended Data Fig. 8) and by inspecting the residuals and prediction uncertainty. The model attained an OOB r^2^ of 0.6 with a OOB prediction error (MSE) of 0.015 (Extended Data Fig. 8). We checked for the presence of spatial correlation in the residuals in several ways. To quantify the spatial proximity between geographical units, we used a topological approach that considers geographical units as neighbors when they share a contiguous boundary. This allowed us determining which areal units should be considered as neighbors, taking into account natural barriers and the non-convex shape of the continent. We computed the graph of neighborhood based on topology and used it to test autocorrelation with global and local Moran tests^75^. Both tests were not significant, indicating the absence of spatial patterns in the residuals of the model (Extended Data Fig. 9). Finally we mapped the prediction uncertainty (Extended Data Fig. 10) and found that it was generally below 20%. We used the jackknife-after-bootstrap^76^ for bagging to estimate the standard errors based on out-of-bag (OOB) predictions with the package ranger^77^.

### Variable importance

To hierarchize the effect of the potential drivers of RASR, we used a permutation variable importance approach with the ranger package^77^. This approach considers a variable important if it has a positive effect on the prediction performance. To evaluate this, a tree is grown in the first step, and the prediction accuracy in the OOB observations is calculated. In the second step, any association between the variable of interest X_i and the outcome is broken by permuting the values of all individuals for X_i, and the prediction accuracy is computed again. The difference between the two accuracy values is the permutation importance for X_i from a single tree. The average of all tree importance values in a random forest then gives the random forest permutation importance of this variable. The procedure is repeated for all variables of interest.

### Influence of variables on RASR

We explored the influence of the 12 environmental variables on RASR in two steps.

First, we analyzed the influence of the variables separately with centered Individual Conditional Expectation Plots^78^ (c-ICE Plots). This type of plot shows the effect of a given predictor X_i on the variable of interest Y (RASR here) for an average situation, meaning for all other covariates at their average value, like in Partial Dependent Plots, and also for each individual environmental condition – meaning all observed combinations of the other covariates. To achieve this, it computes the expected values of Y across the range of the covariate X_i, while keeping all other covariates constant. C-ICE plots thus allows us to see the average effect of the covariates, and also the heterogeneity of this effect in individual combinations of covariates across the predictors’ space. In these plots, the expected values are centered to their value at the lowest value of the covariate X_i in order to facilitate the comparison among individual conditions.

In a second step, we explored potential interactions between predictors by a double approach. We started by drawing all pairwise 2-D Partial dependent plots, restricting the co-variables to lie within with convex hull of their possible values, and searched for potential interactions. Then we used the joint-Variable Importance Measure (JVIMP) test^79^ to assess the significant interactions. In this approach, two co-variates are paired and their paired VIMP is calculated. The VIMP for each separate variable is also calculated. The sum of these two values is referred to as ’Additive’ importance. A large positive or negative difference between ’Paired’ and ’Additive’ indicates an association worth pursuing if the univariate VIMP for each of the paired-variables is reasonably large. The find.interaction () function of the R package randomForestSRC^80^ was used with the option vimp. We found no evidence of significant interaction.

### Comparison with the invasion patterns in other groups

We compared our results on earthworm to data of invasion patterns on several aboveground organisms. These data came from a global database of invasive species^4^. We restricted our analysis on taxa with the best geographical coverage across North America: mammals, birds, plants, and spiders. This database reports the number of invasive species in the TGWD-4^4^ geographical units previously mentioned. We computed the number of alien earthworm species in the TDWG-level 4 units from the complete EWINA database, excluding Greenland, (Extended Data Fig. 6) and compared it to the above ground organisms with a Spearman correlation.

### Data availability

Data are available online on the Zenodo datawarehouse. EWINA_RICH, the database of earthworm native and alien species richness in North America at the two spatial resolutions, together with the aboveground alien species richness in TDWG4 units, is available at https://doi.org/10.5281/zenodo.6421014. EWINA_IPATHS, the database of intercepted earthworm introduction pathways from 1945 to 1975 is available at https://doi.org/10.5281/zenodo.6408609. EWINA_SP, the tabular data of Ecological profile of earthworm species found in North America, including geographical origin, alien status, functional role and reproduction mode is available at https://doi.org/10.5281/zenodo.6462831. EWINA_1st_RECORDS, the list of first year of observation of each earthworm species in North America can be found at https://zenodo.org/record/6759725. The sources of the environmental data are given in Extended Data Table 1. The list of data sources is provided in Supplementary Information.

### Code availability

R scripts used for the analyses are available at https://github.com/JeromeMathieuEcology/GlobalWorming

## Acknowledgments

We thank the people who helped with digitizing information on the distribution of native and alien earthworm species, particularly E. Pedarros, S. Dandrifosse, C. Mathieu, J. Nabias, and T. Allain. We highly appreciate the comments and suggestions of Isabelle Gounand, Gérard Lacroix and Olivier Moine, and the help of W. Dawson and R. Early regarding the use of their databases, as well as the help of Marvin Wright regarding the use of the package ranger. Finally, we thank the Animal Plant Health Inspection Service from the United States Department of Agriculture and the Invasive Alien Species & Domestic Programs Section from the Canadian Food Inspection Agency for their help regarding the import-export regulation laws. Funding was provided by the France-Stanford Center for Interdisciplinary Studies.

## Author contributions

J.M. and E.H. conceived the original idea and secured the principal funding. J.M. curated the data and ran the analyses. J.M. J.W.R. and C.F. searched for data, J.M. wrote the first draft and all authors reviewed and edited the manuscript.

## Additional information

**Supplementary Information** is available for this paper.

**Correspondence and requests for materials** should be addressed to Jérôme Mathieu

**Reprints and permissions information** is available at www.nature.com/reprints.

## Extended data

**Extended Data Fig. 1.**
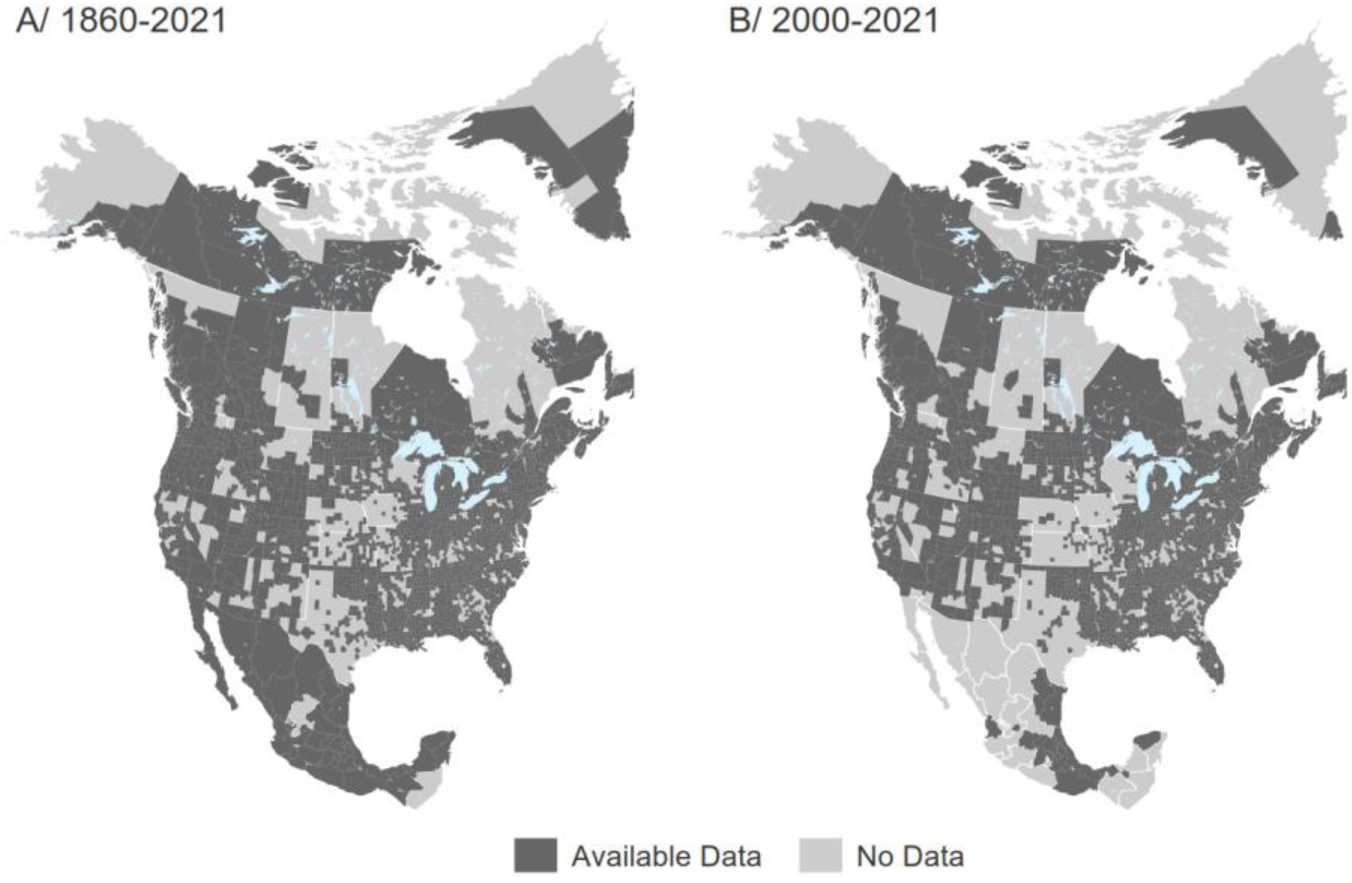
Geographical coverage of the EWINA database. Data that are available in the EWINA database are indicated in dark grey, while geographical units with no data are indicated in light grey. A/ Complete coverage. B/ Coverage for the period 2000-2021. Only the data from 2000-2021 were used for modelling RASR.

**Extended Data Fig. 2.**
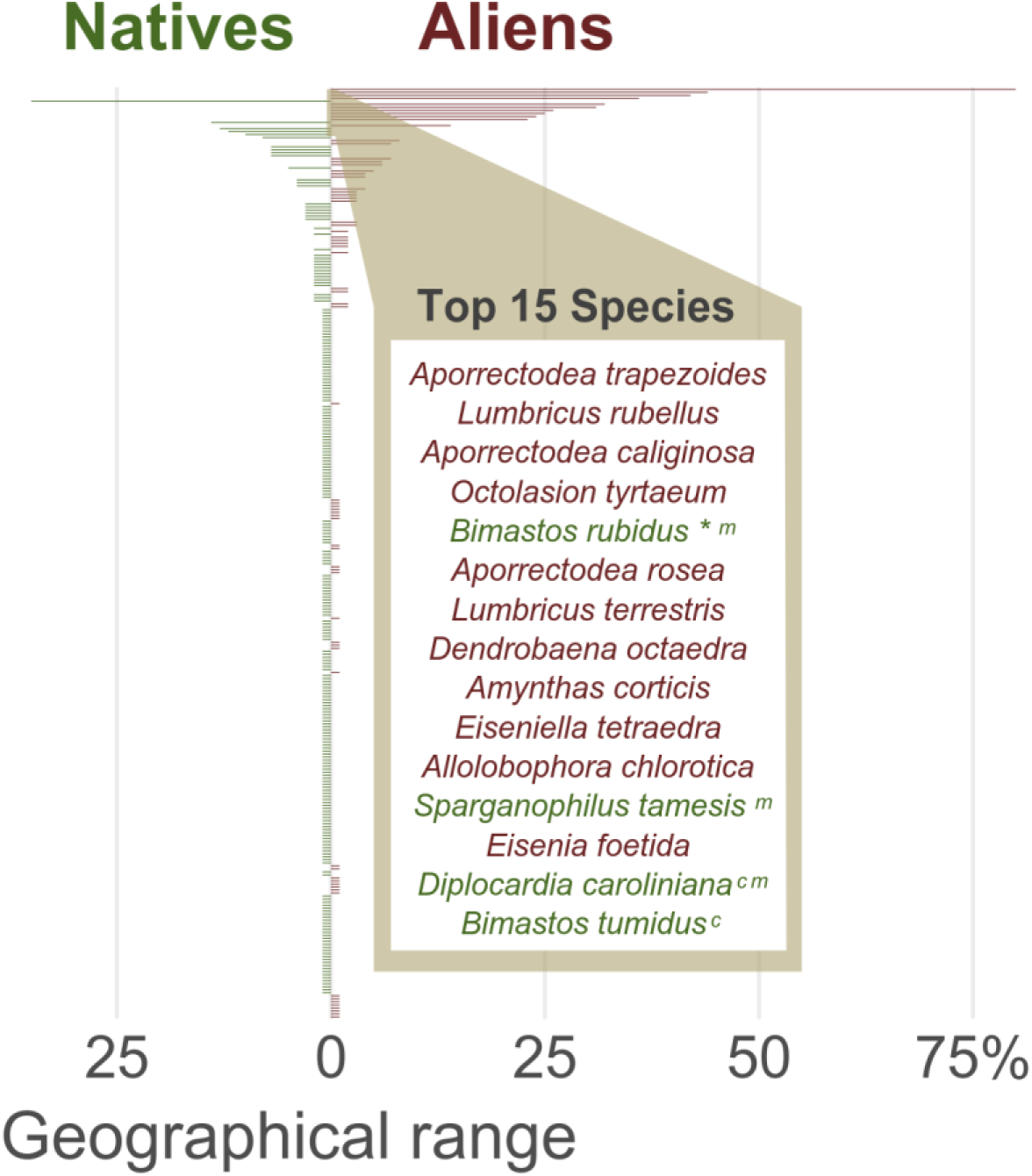
Geographical range of Native and Alien species across North America. Each horizontal bar represents the geographical range of a species. The range is expressed as the proportion of geographic units where the species occur. The color of the bars indicates the origin of the species. The 15 most widespread species are list in the box, by decreasing order of geographical range. * *B. rubidus* is a synonym of *Dendrodrilus rubidus*, ^m^ : species considered as alien in Mexico, ^c^ species considered as alien in Canada, ^cm^ species considered as alien in Canada and Mexico.

**Extended Data Fig. 3.**
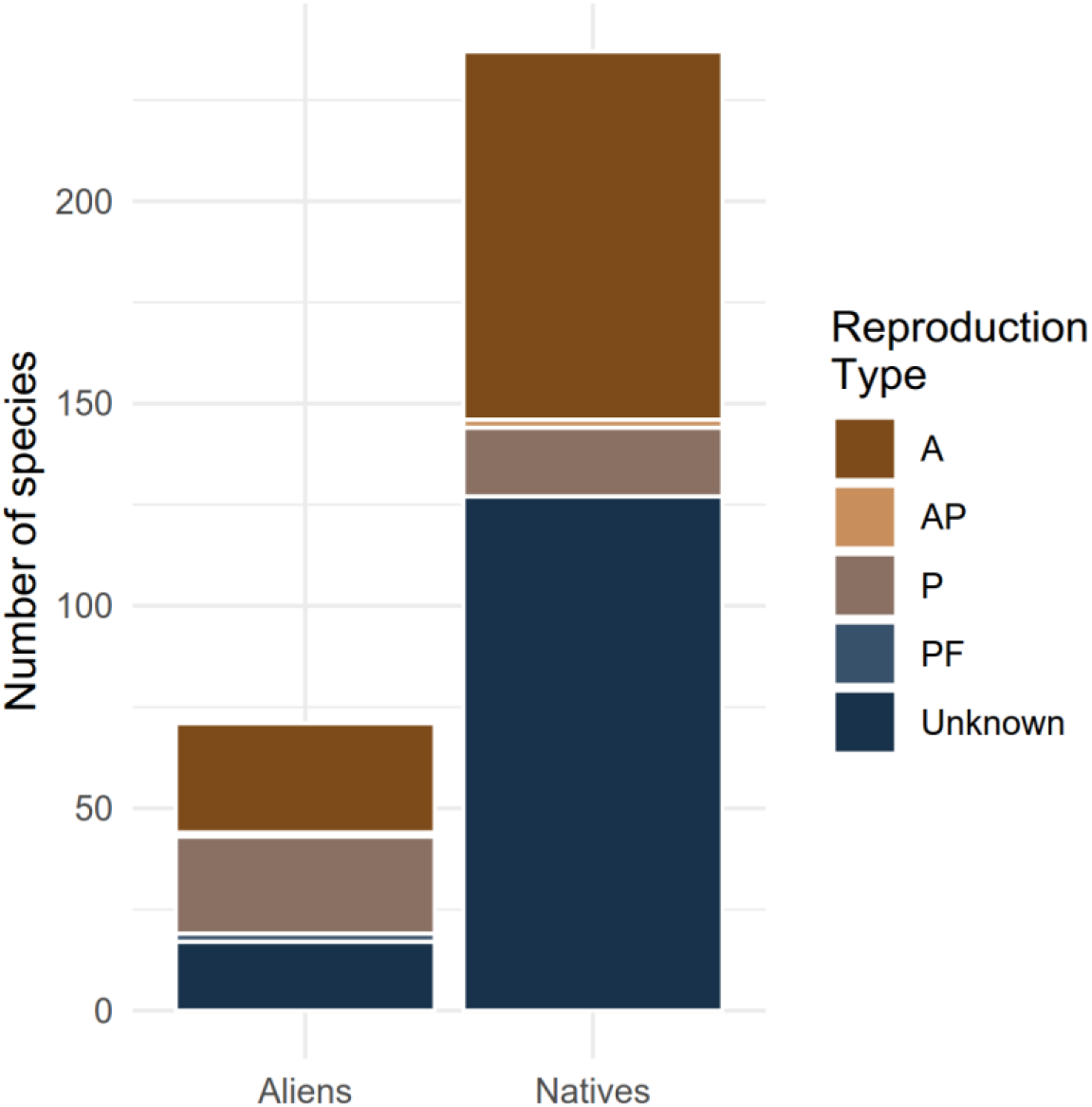
Reproduction type of native and alien earthworm species in North America. A : Amphimictic : reproduction sexual and biparenthal, AP : generaly amphimictic with parthenogenesis in some morphs, P : parthenogenetic, reproduction uniparental, PF : parthenogenesis facultative (χ^2^ = 42.8, p = 1.15 10^-8^).

**Extended Data Fig. 4.**
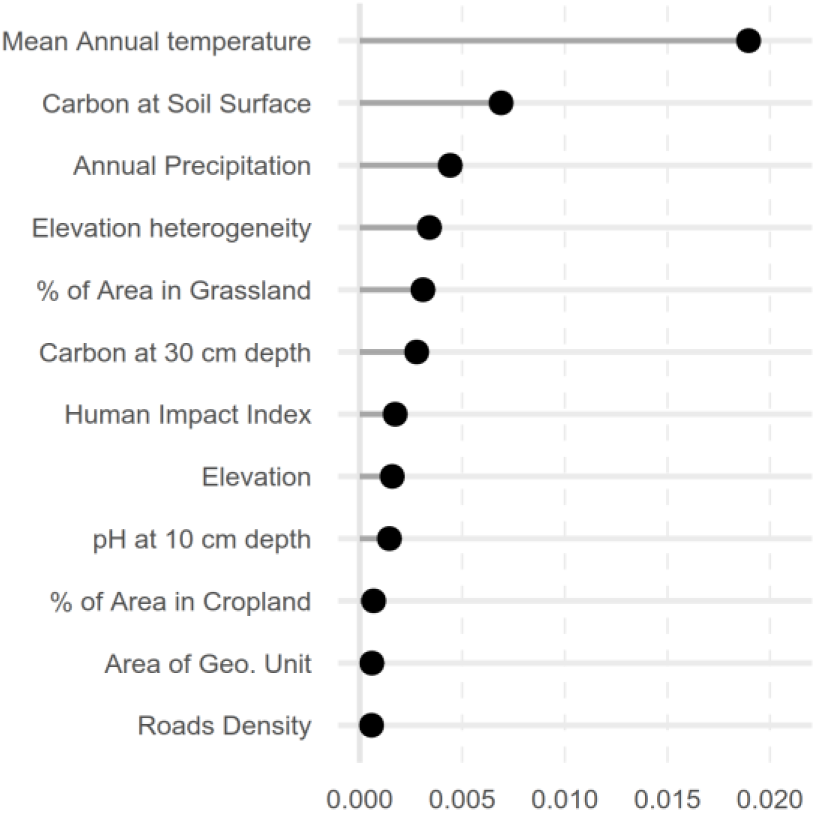
Variable importance in the random forest model of earthworms’ RASR. Variable importance represents the magnitude of accuracy loss of the predictions across all trees when randomizing the considered variable. The procedure is repeated for all variables of interest. Variable importance was computed with the ranger package^77^.

**Extended Data Fig. 5.**
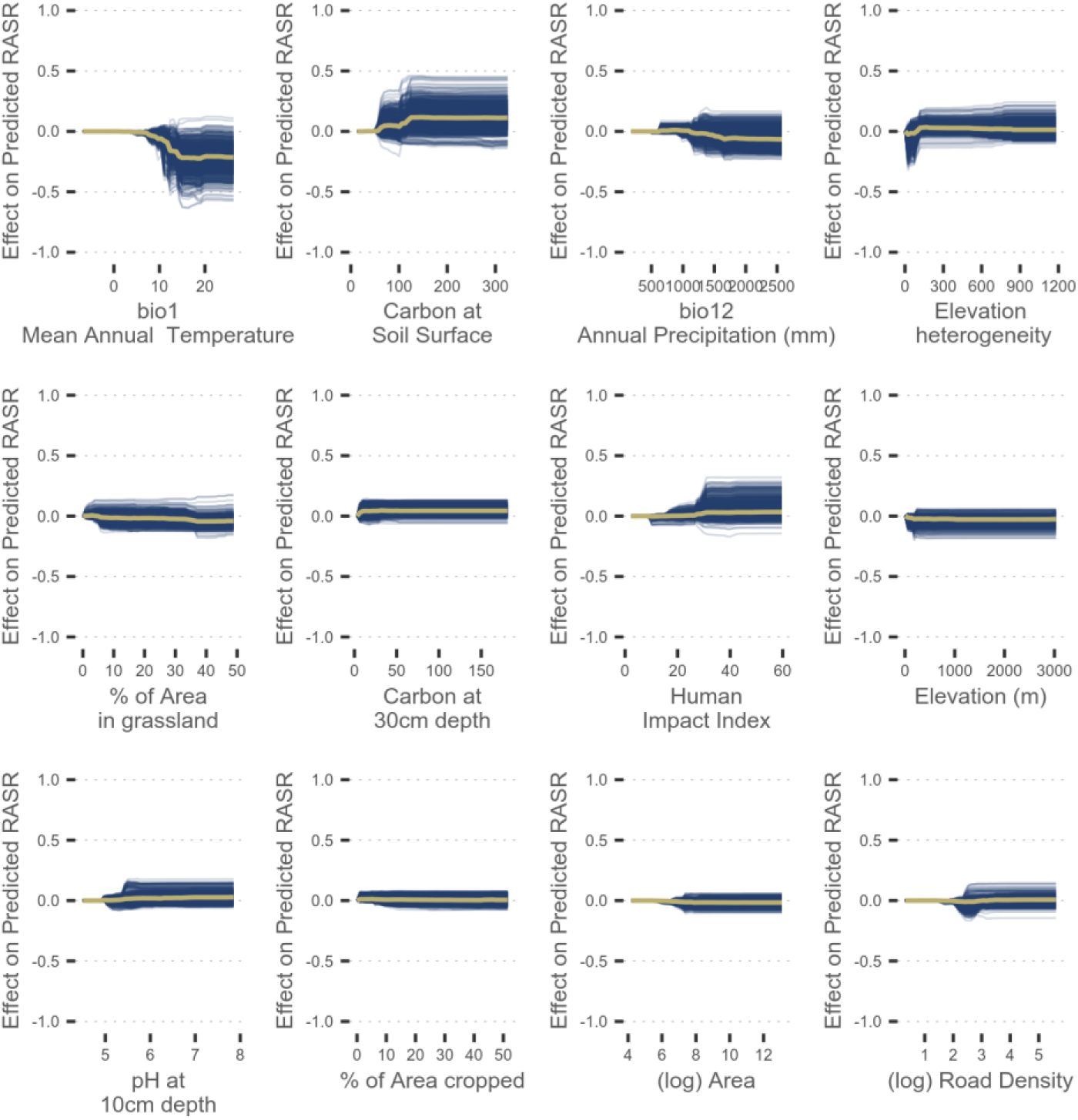
C-ICE plots: influence of the covariates on predicted RASR. The thick yellow line indicates the effect of the covariate for an average environmental condition (All other covariates being at their average value). The blue lines indicate the effect of the covariate for the individual condititions across the training dataset. The heterogeneity of the individual blue lines indicates the variability of the predictor across individual conditions.

**Extended Data Fig. 6.**
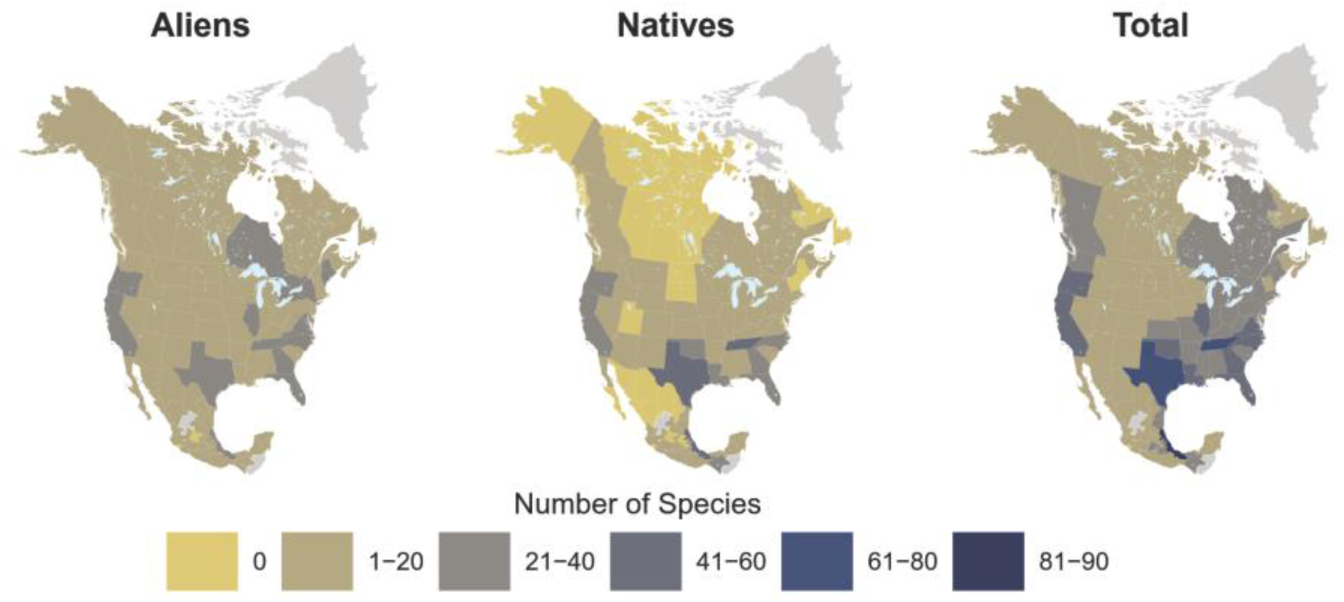
Earthworm species richness in the TDWG-level 4 geographical units. Maps of the species richness of earthworm across the TDWG-level 4 regions. The TDWG regions are standardized geographical units defined to compare species richness across taxa at large scale ^31^. The maps detail the (A) Alien, (B) Native, and (C) Total species richness of earthworm. These data were used for the comparison with above ground taxa (Fig. 2).

**Extended Data Fig. 7.**
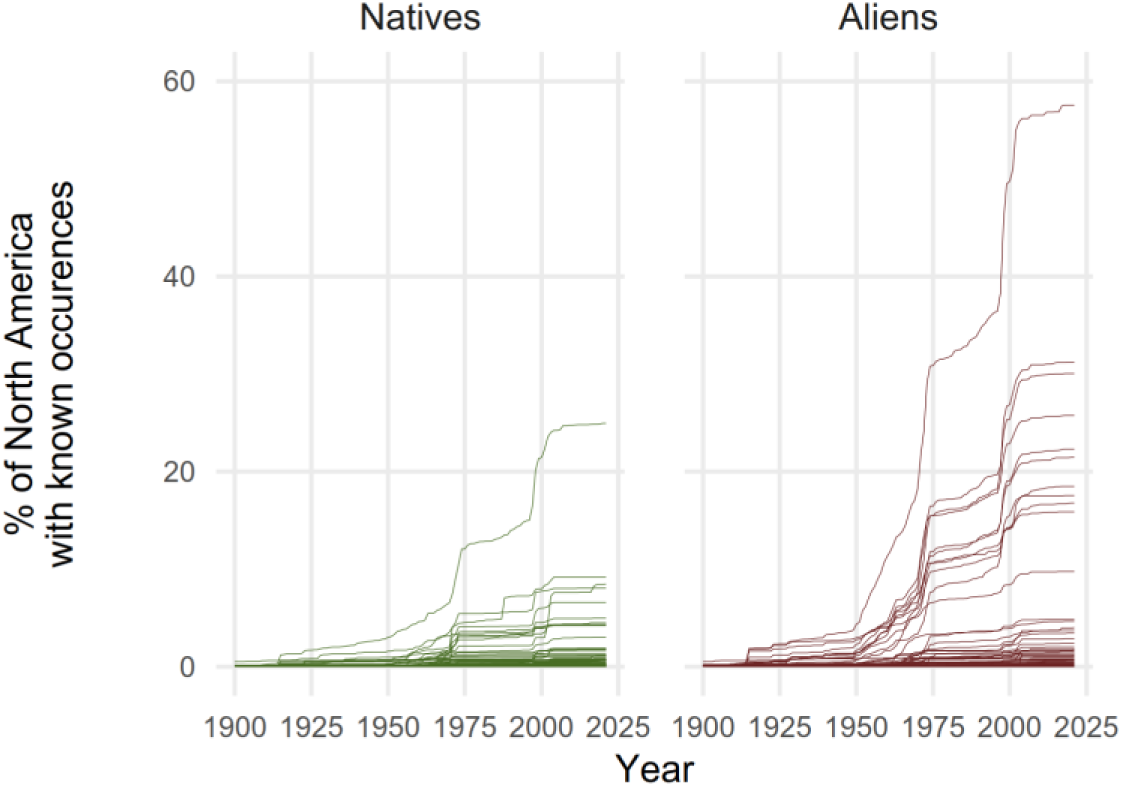
Temporal dynamics of the known geographical range of native and alien earthworm species across North America. Each line represents the geographical range of a species over time. Geographical range is expressed as the percentage of geographical units (county level, Extended Data Fig. 1) that are occupied by a species.

**Extended Data Fig. 8.**
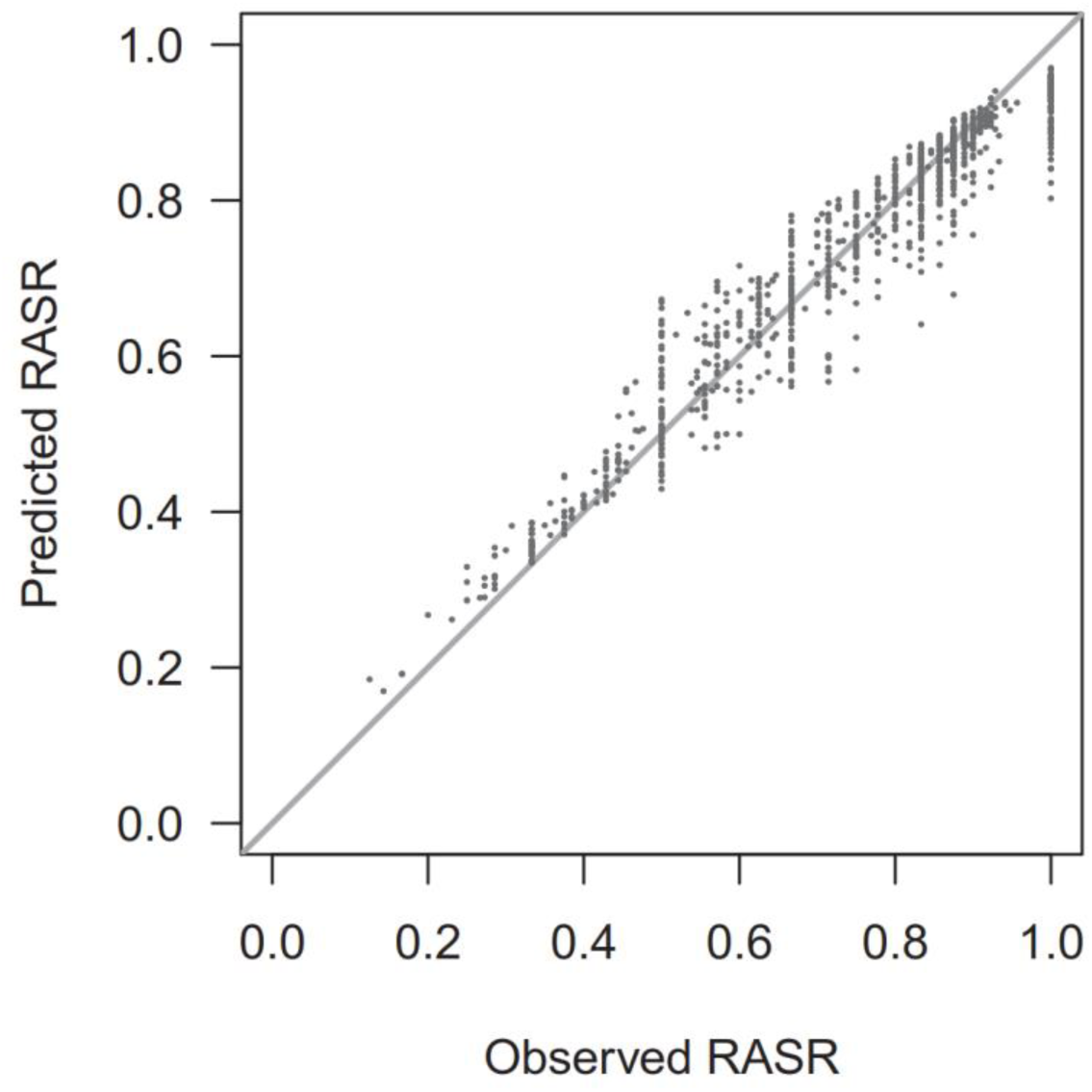
Model performance. This plot shows the predicted value of each observation, using a model in which the observation was not used to train the model (OOB test data), against the observed RASR. The closer the data are from the line, the best the model performs. We can see that the model tends to slightly underestimate RASR in actual high values RASR and slightly overestimate RASR values in actual low values of RASR. OOB r^2^ is 0.6.

**Extended Data Fig. 9.**
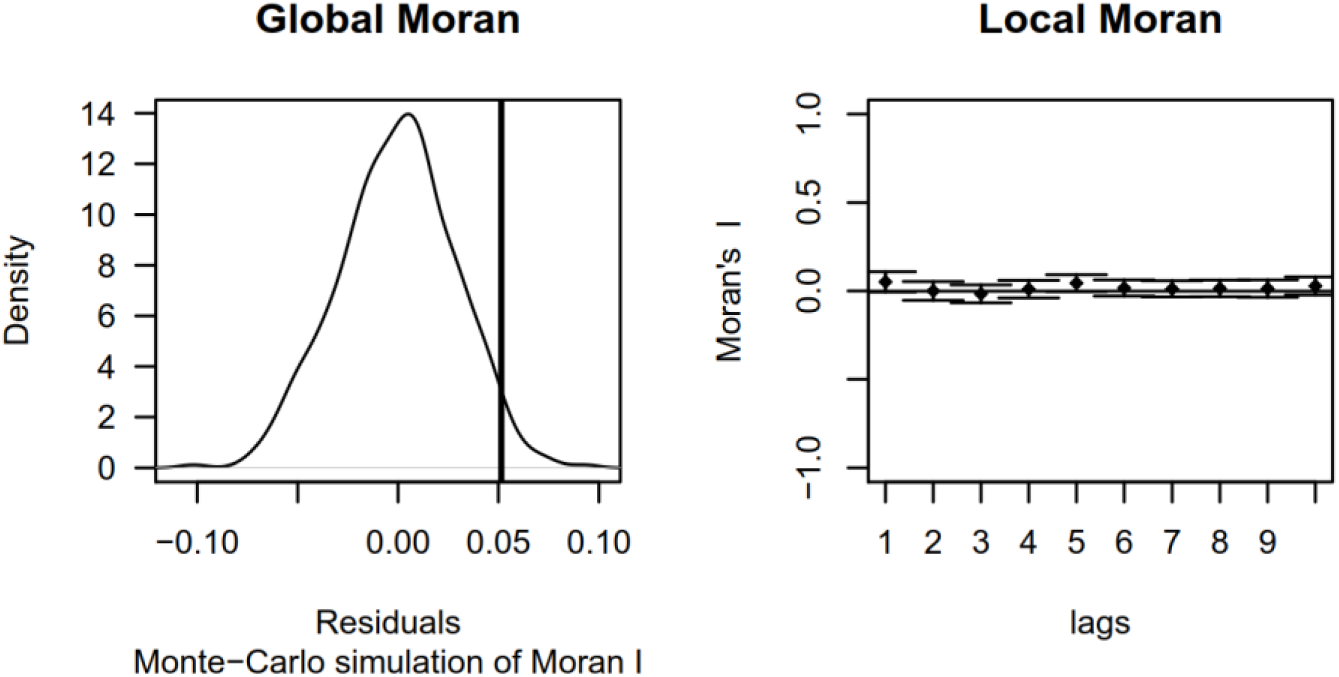
Test of the autocorrelation in the residuals. These plots are aimed at testing the existence of spatial correlation in the residuals of the random forest. (A) The global Moran’s I test the existence of global correlation by a permutation procedure. The distribution of the random values are indicated by the bell shaped curve. The observed value of the global Moran, indicated by the vertical bar, falls within the null distribution, indicating an absence of autocorrelation. (B) The correlation is tested between geographical units separated by an increasing number of neighbors (lags), indicated on the X axis. This allows to test the existence of spatial correlation at precise lags of neighboring. The horizontal line indicates a null spatial correlation. The whiskers indicate the confidence interval of the Moran indices for each lag. There is significant correlation when the whiskers do not intercept the horizontal line. Here, the trend is fairly flat with the majority of Moran’s value being near 0, indicating the absence of spatial autocorrelation of the residuals.

**Extended Data Fig. 10.**
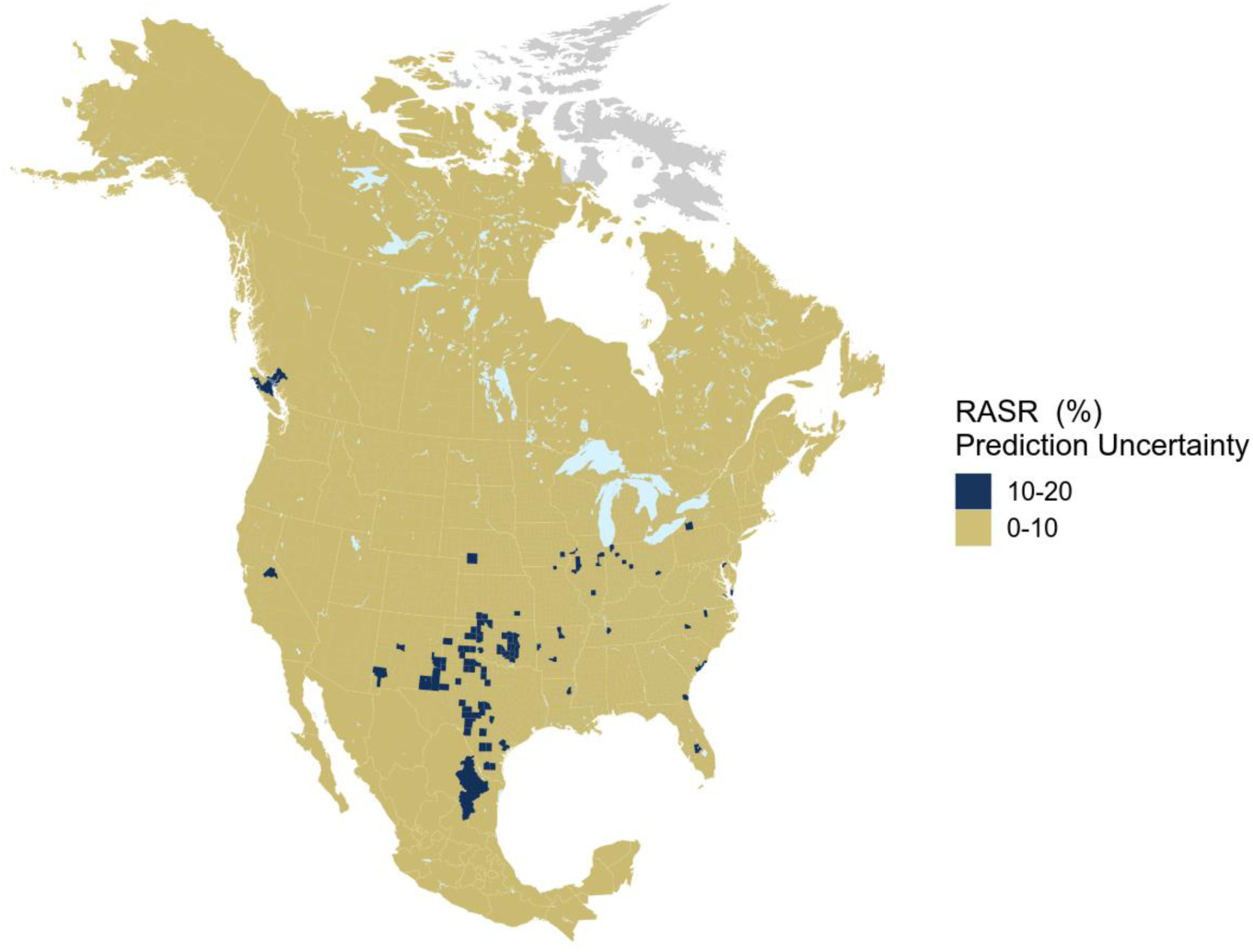
Map of predictions’ uncertainty. This map shows the uncertainty of the predictions made by the model. This is useful to interpret the predicted values mapped in Fig.1. We see that the uncertainty is overall relatively low.

**Extended Data Table 1.** Detail of the environmental covariates used to model earthworm RASR.

|  | Variable | Source | Weblink | Native Spatial Resolution |
| --- | --- | --- | --- | --- |
| Geographical features | Area | GADM 3.6 | https://www.gadm.org/data.html | - |
|  | Elevation | gtopo30 (USGS) | https://doi.org/10.5066/F7DF6PQS | 30 Seconds |
|  | Elevation heterogeneity |  |  |  |
| Climate | bio1 - Annual Mean Temperature | Worldclim | https://www.worldclim.org/ | 10 minutes |
|  | bio12 - Annual Precipitation |  |  |  |
| Human Activity | Human Impact Index | SEDAC | https://sedac.ciesin.columbia.edu/data/set/wildareas-v2-human-influence-index-geographic/data-download | 30 Seconds |
|  | % of Area in grassland | HYDE 3.2.1 | https://easy.dans.knaw.nl/ui/datasets/id/easy-dataset:74467 | 5 Minutes |
|  | % of Area in cropland |  |  |  |
|  | Roads Density | Natural Earth | https://www.naturalearthdata.com/downloads/10m-physical-vectors/ | 1:10m |
| Soil Properties | Soil pH at 10 cm depth | SoilGrids | https://soilgrids.org/ | 1 km |
|  | C0 - Soil Carbon Content on Soil Surface |  |  |  |
|  | C30 - Soil Carbon Content at 30 cm depth |  |  |  |

